# Promoting translational readthrough to augment fibrillin-1 (FBN1) deposition in Marfan syndrome fibroblasts: A proof-of-concept study

**DOI:** 10.1101/2022.11.23.517642

**Authors:** Zerina Balic, Dirk Hubmacher

## Abstract

Marfan syndrome (MFS) is a connective tissue disorder characterized by long bone overgrowth, enlargement of the aorta, ocular anomalies and other symptoms. Current treatment focuses on managing aortic aneurysms to avoid dissection or rupture. However, no cures are available. MFS is caused by one of >1,800 dominant pathogenic variants in *FBN1*, which encodes the extracellular matrix (ECM) protein fibrillin-1. A significant number of *FBN1* variants result in premature termination codons (PTCs). Recently, small molecules were identified that can promote translational readthrough of PTCs and were evaluated in preclinical and clinical trials for several genetic disorders. Here, we show that the translational readthrough drugs ataluren and gentamicin ameliorated FBN1 deposition in some MFS patient-derived skin fibroblast lines harboring PTC variants in *FBN1*. In contrast, inhibitors of NMD were cytotoxic to the skin fibroblast lines that we analyzed. We conclude that promoting translational readthrough of PTC variants in *FBN1* could result in a therapeutic benefit for MFS patients with specific PTCs in *FBN1* and that its efficacy will likely depend on the PTC sequence context, the amino acids that are incorporated in FBN1 after PTC suppression and the overall increase of FBN1 deposition in the ECM.

## Introduction

Premature termination codons (PTCs) arise from single nucleotide polymorphisms in the coding region of a gene, where they change the codon for an amino acid into one of three stop codons, i.e. TAA, TAG, or TGA. PTCs account for an estimated 11% of all pathogenic variants that cause inherited human genetic disorders [1]. The presence of a PTC can elicit nonsense-mediated mRNA decay (NMD), resulting in a marked reduction of mRNA, or lead to premature termination of mRNA translation, resulting in the synthesis of a truncated and potentially non-functional or even toxic protein product [2, 3]. However, an estimated 5 – 25% of mRNA transcripts carrying a PTC can escape NMD [4, 5]. Because PTCs are read through at a higher level (up to 1%) compared to normal stop codons (0.001 – 0.1%), further promoting translational readthrough with small molecules with or without NMD inhibition represents a potential strategy to augment the amount of functional full-length protein in patients with genetic disorders caused by PTCs [6, 7]. Since the mid-1980s, several drugs were identified that promoted translational readthrough in vitro, in patient-derived cells and in mouse models, some of which have entered clinical trials [8, 9]. It has been shown that aminoglycoside antibiotics such as G418 or gentamicin can promote translational readthrough of PTCs resulting in increased amounts of full-length protein without substantially altering the efficiency of the natural termination codon [10]. Subsequently, translational readthrough promoting small molecules have been evaluated in preclinical and clinical trials targeting genetic disorders including lysosomal storage disorders, hemophilia, muscular dystrophies or cystic fibrosis with somewhat variable outcomes [8, 9, 11–13]. In these studies, the outcome was largely determined by the threshold amount of functional protein required to ameliorate disease progression. Translational readthrough efficiency depends on the sequence context of the PTC, the specific readthrough drug and the amount of mRNA that escapes NMD [14]. In dominant disorders, the identity of the inserted amino acid, which is determined by the PTC (UAA and UAG can be recognized by glutamine, tyrosine or lysine tRNAs, UGA by tryptophan, arginine or cysteine tRNAs) is an important factor in gauging their efficacy due to potential dominant negative effects of the variant allele on the wild-type allele [7, 15]. Decoding of the PTC is guided by the shape of the base pairs in the 1 or 3 position, where UAG preferred mismatches in the 1 position, UGA in the 3 position and UAA showed no preferences [7].

Marfan syndrome (MFS) is a connective tissue disorder where musculoskeletal, cardiovascular, pulmonary, and ocular tissues are compromised [16, 17]. Thoracic aortic aneurysm is the most severe symptom in MFS since it can lead to life-threatening aortic dissections and potentially ruptures [18]. MFS is caused by one of 1,847 dominant pathogenic variants in fibrillin-1 (*FBN1*) and has a prevalence of 2-3 in 10,000 newborns [19]. Fibrillin-1 forms the core of fibrillin microfibrils in the ECM that serve as an ECM scaffold and regulate TGF⍰ and BMP growth factors [20–22]. The current standard of care for patients with MFS includes pharmacological blood pressure control to reduce the stress on the aortic wall, preventative surgical replacement of dilated sections of the aorta, lens replacement, and physiotherapy, surgery or pain management to alleviate musculoskeletal symptoms [23]. Dominant *FBN1* variants can result in haploinsufficiency, where the amount of normal fibrillin-1 microfibrils in tissues is reduced, or dominant negative effects, where FBN1 protein produced from the variant allele suppresses the function of wild-type FBN1 [24–29].

Since a significant number of pathogenic *FBN1* variants result in PTCs, promoting translational readthrough could be a potential therapeutic strategy for MFS by increasing the amount of functional FBN1 microfibrils in the ECM. On the other hand, promoting the synthesis of a FBN1 protein that may now carry an altered amino acid residue in the position of the PTC could have unwanted dominant negative effects. Here, we present a proof-of-concept study that promoting translational readthrough in MFS patient-derived fibroblasts harboring a PTC in *FBN1* could in principle augment FBN1 deposition in the ECM. We provide an *in-silico* analysis of the *FBN1* PTCs and consider the size of the potential patient population. Using MFS patient-derived fibroblasts, we demonstrate that some, but not all, cell lines responded to treatment with translational readthrough drugs with enhanced deposition of FBN1 microfibrils. Finally, we provide a possible decision tree that would stratify MFS-causing PTCs in order to identify individuals that could benefit from treatment with a translational readthrough drug and outline the required preclinical tests in a personalized medicine approach.

## Materials and Methods

### Cell culture and drug treatments

Primary MFS fibroblast cell lines were characterized previously and purchased from the Coriell Institute for Medical Research (Camden, NJ) as live cultures [30] (Table 2). MFS fibroblasts were cultured in Dulbecco’s modified Eagle’s medium (DMEM), supplemented with 10% fetal bovine serum (FBS), 1% L-glutamine, 100 U/ml penicillin and 100 ⍰g/ml streptomycin (complete DMEM). Fibroblasts were cultured in a 5% CO_2_ atmosphere in a humidified incubator at 37 °C. Cells were passaged with trypsin-EDTA in a 1:3 – 1:5 ratio when MFS fibroblasts reached confluency. The small molecules promoting translational readthrough or inhibiting NMD used in this study and their respective solvents and sources are listed in Table 3. Small molecules were dissolved as stock solutions and frozen in single-use aliquots. Working solutions of the small molecules were prepared by diluting stock solutions in complete DMEM prior to addition to the MFS fibroblasts. The largest volume of solvent was diluted in complete DMEM and used as control for the individual experiments. For immunostaining, 50,000 cells per chamber were seeded in 8-well chamber slides (Celltreat Scientific Products, Pepperell, MA). Drug treatment was initiated at day 1 after cell seeding and a fresh aliquot was added at day 3 after cell seeding. MFS fibroblasts were processed for immunostaining at day 5 after cell seeding. To collect conditioned medium and cell lysates, MFS fibroblasts were seeded in 6-well plates at 150,000 cells per well and treated with 20 ⍰M ataluren or 1 mM gentamicin at day 1 (in 2 ml complete DMEM) and day 3 (in 1.5 ml serum-free complete DMEM) after cell seeding. Conditioned medium and cell lysate were collected at day 5 after cell seeding and processed for western blot analyses.

### Immunofluorescence staining

Cell culture medium was removed and the MFS fibroblast layer was rinsed with phosphate-buffered saline (PBS) and fixed for 5 min with ice-cold 70% methanol/30% acetone (Thermo-Fisher, Waltham, MA). Cells were rinsed with PBS and non-specific antibody binding sites were blocked with 10% normal goat serum (Jackson ImmunoResearch Laboratories, West Grove, PA) diluted in PBS (blocking buffer) for 1 h at room temperature (RT). Cell layers were incubated with primary antibodies against fibrillin-1 (described previously, 1:1000) and fibronectin (clone FN-15, 1:1000, Millipore-Sigma) diluted in blocking buffer for 2 h at RT [31]. Cells were rinsed 3 × 5 min with PBS and incubated with AlexaFluor-labeled secondary goat-anti-mouse or goat-anti-rabbit antibodies (1:350, Jackson ImmunoResearch Laboratories, West Grove, PA) diluted in blocking buffer for 2 h at RT. Cells were rinsed 3 × 5 min with PBS and mounted using ProLong Gold Antifade Reagent containing 4′,6-diamidino-2-phenylindole (DAPI) to stain cell nuclei (Thermo-Fisher, Waltham, MA). Cells were imaged using a Zeiss Axio Observer Z1 Fluorescence motorized microscope. For quantification, images from 3-4 fields of view were captured. To allow comparison of experimental conditions, exposure times were kept constant and adjusted to the strongest signal for each cell line and experimental condition. Image analysis was performed with Zeiss Zen Microscope Software and the arithmetic mean pixel intensity was used for quantitative analysis. The fluorescence signal was quantified as mean pixel intensity and normalized to DAPI signal. Statistical analysis was performed using one-way ANOVA with a significance level of p <0.05 to compare treatment groups. Statistical comparison between individual samples was done post-hoc using the Tukey’s Multiple Comparison test where a p-value <0.05 was considered significant using OriginPro 2019 software (OriginLab, Northampton, MA).

### Western blot analysis

Conditioned medium was cleared by centrifugation for 5 min at 14,000 rpm. Proteins in 56 μl conditioned medium were denatured with by adding 14 μl reducing SDS-polyacrylamide gel electrophoresis (PAGE) loading buffer (0.5 M Tris pH 6.8, 5% 2-mercaptoethanol, 20% SDS, 1% Bromophenol blue, 50% Glycerol) and incubation for 5 min at 95 °C. The cell layer was rinsed with PBS and lysed in 100 ⍰l of 0.1% NP40, 0.01% SDS, 0.05% Na-deoxycholate in PBS for 5 min on ice. Cell lysates was cleared by centrifugation at 14,000 rpm for 15 min at 4 °C. Proteins in the supernatant were denatured by adding 14 μl reducing SDS-PAGE loading buffer to 56 μl lysate followed by incubation at 95 °C for 5 min. Proteins in 65 μl of conditioned media and cell lysates per lane were separated on 6% polyacrylamide gels using SDS-polyacrylamide gel electrophoresis (SDS-PAGE) and transferred on a poly-vinylidene difluoride (PVDF) membrane (Immobilon-FL, Merck Millipore Ltd., Burlington, MA) using the MiniProteas Blotting Module (Bio-Rad, Hercules, CA) for 1.5 h at 70 V with 25 mM Tris, 192 mM glycine, 20% methanol as transfer buffer. Upon completion of the transfer, membranes were blocked with 5% (w/v) milk in 10 mM Tris-HCl, pH7.2, 150 mM NaCl (TBS) for 1 h at RT and incubated with primary antibodies against FBN1 (1:500), fibronectin (1:500) or GAPDH (MAB374, 1:4000, Millipore-Sigma) diluted in 5% (w/v) milk in TBS including 0.1% Tween 20 (TBS-T) overnight at 4 °C. Membranes were rinsed 3 × 5 min with TBS-T and incubated with IRDye goat-anti-mouse or goat-anti-rabbit secondary antibodies (1:10,000 Jackson ImmunoResearch Laboratories, West Grove, PA) diluted in 5% (w/v) milk in TBS-T, for 2 h at RT. Membranes were washed 3 × 5 min with TBS-T, once with TBS and imaged using a Licor Imaging system. Fluorescent intensities of individual bands representing FBN1 and fibronectin were quantified using ImageJ Fiji.

## Results

### In silico analysis of MFS-causing PTC variants in *FBN1*

The 2,871 amino acids of FBN1 are encoded by 8,613 nucleotides distributed over 65 coding exons of the *FBN1* gene. As of July 2022, the Universal Mutation Database (UMD-FBN1) listed 1,847 different pathogenic *FBN1* variants identified in 3,077 samples [19]. The majority of *FBN1* variants are single nucleotide variants (1,225; 66.3%) resulting in missense variants (1,015; 55%) or in PTCs (210; 11.4%) (Fig 1A). The *FBN1* variants causing classical MFS are distributed over the entire FBN1 protein with very little genotype-to-phenotype correlation (Fig 1B, top). Exceptions are a region spanning exons 24 – 32, where pathogenic variants can cause severe and early onset neonatal MFS, and exons 64 – 65, where pathogenic variants can cause neonatal progeroid syndrome, which resembles MFS but includes severe lipodystrophy [24, 32–34]. In addition, pathogenic *FBN1* variants in the TGFβ-binding protein-like (TB)/8-Cys domain 5 can cause acromicric or geleophysic dysplasia, which present as “opposite” to MFS and are collectively characterized by short stature, joint contractures, tight skin, and enlarged skeletal muscles [35, 36]. When mapping the distribution of PTC variants in *FBN1*, we observed that exon 7 and 10, which encode the calcium-binding epidermal growth factor (cbEGF) domain 1 and the proline-rich domain, respectively, harbored the highest percentage of PTC variants (87% and 66.7%, respectively) and only very few missense variants (4.3% and 6.6%, respectively) (Fig 1B, bottom). In contrast, exons 22, 42 and 52 encoding the C-terminal part of hybrid domain 2, the C-terminal part of TB/8-Cys domain 5, and cbEGF 32, respectively, were devoid of PTCs. In addition, four clusters of PTC variants were identified, where the average percentage of PTC variants per exon was higher than 25%. These clusters include *FBN1* exons 7 – 10 (except for exon 8), 36 – 39, 53 – 58 (except for exon 55) and 64 – 65. Of note, PTCs located in the penultimate and ultimate *FBN1* exons (64 – 65) are likely to escape NMD and may result in a truncated FBN1 protein [37].

**Fig 1.**
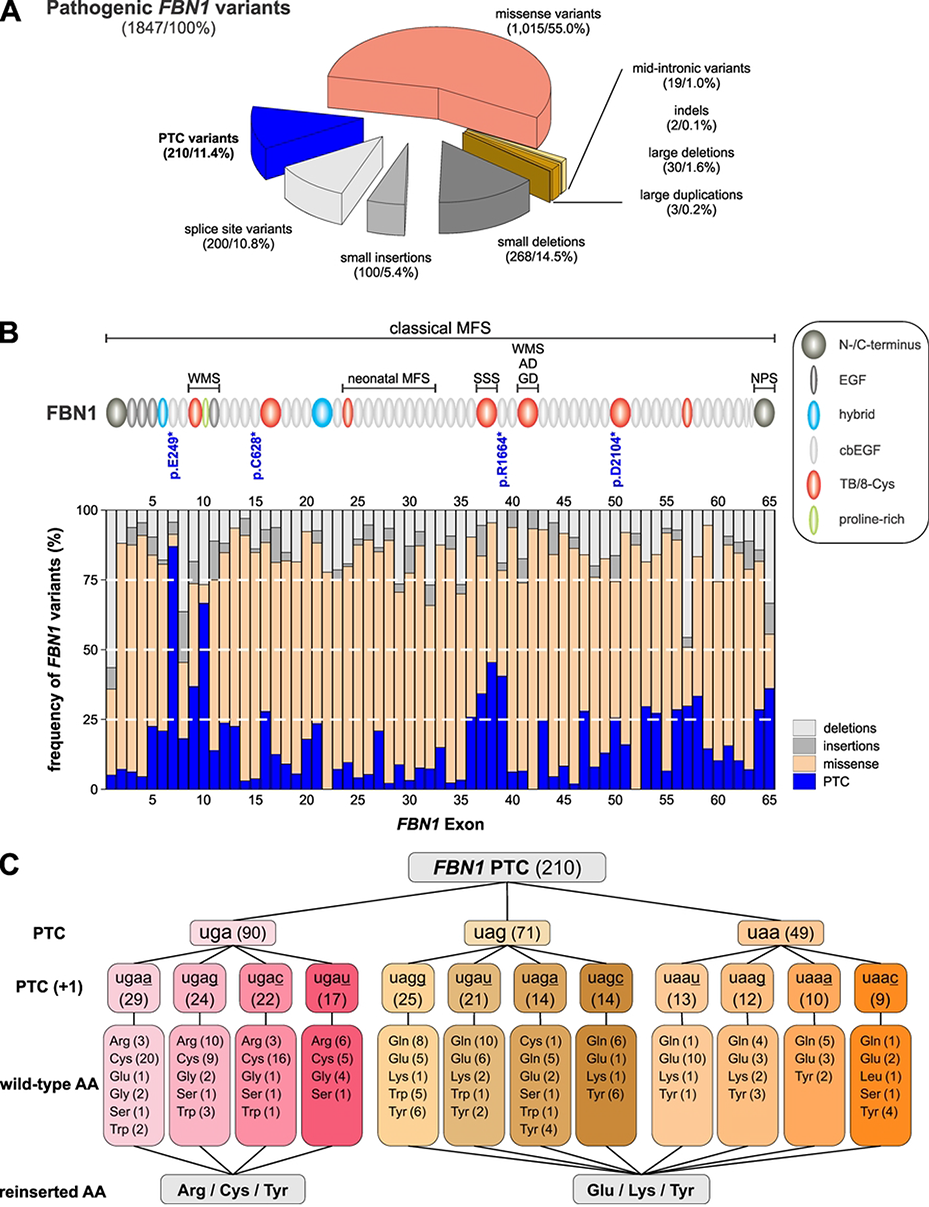
*In silico* analysis of MFS-causing PTCs in *FBN1*. **(A)** Frequency of pathogenic *FBN1* variants based on the UMD-*FBN1* database [19]. The absolute number of *FBN1* variants and their relative proportions are indicated in brackets. Small insertions/deletions are defined as affecting less than one exon and large deletions and duplications as affecting more than one exon; **(B)** Percentage of PTC variants in each *FBN1* exon compared to other types of variants. The cartoon on top depicts the domain organization of FBN1 and correlates individual exons with the FBN1 domains. Similar sized FBN1 domains, such as TB/8-Cys domains appear distorted since they can be encoded by one or two *FBN1* exons. The neonatal region and exons where variants can cause neonatal progeroid syndrome (NPS) are indicated [34, 38]. Domains where *FBN1* variants can cause acromelic dysplasias (Weill-Marchesani syndrome, WMS; acromicric dysplasia, AD; geleophysic dysplasia, GD) or stiff skin syndrome (SSS) are indicated [35, 39]. FBN1 protein variant numbering is based on the FBN1 protein sequence NP_000129.3; **(C)** Frequency of individual PTCs in *FBN1*, the nucleotide in the +1 position (underlined) the mutated wild-type amino acids and the amino acids that could be potentially incorporated after PTC suppression.

Since the potential success of promoting translational readthrough depends on the PTC sequence and its context, we determined the frequency for each of the three stop codons (UAA, UAG, UGA) and analyzed their sequence context in *FBN1* (Fig 1C) [40, 41]. The UGA stop codon accounted for >40% of pathogenic PTCs in *FBN1* with an enrichment for A, G, C in the critical +1 position immediately following the PTC. The stop codons UAG (33.8%) and UAA (23.3%) were less frequently represented. While the four possible +1 nucleotides following UAA were almost equally distributed, the +1 nucleotides following UAG were enriched for G and U. UGA PTCs did originate mostly from codons for the amino acids arginine and cysteine, while UGA and UAA originated mostly from codons for glutamine and glutamic acid. Based on the possible amino acids that can be incorporated when promoting translational readthrough, there are 141 of the 210 *FBN1* PTCs where the original amino acid could potentially be reincorporated. In contrast, in 62 of the 210 PTC positions, pathogenic *FBN1* variants that alter the amino acid but do not result in a PTC have been described in individuals with MFS, 37 of which encode cysteine residues in the wild-type *FBN1* sequence.

To estimate the number individuals with MFS due to a *FBN1* PTC variant that could potentially benefit from promoting translational readthrough, we used publicly available demographic data for the US and world population in 2020 and the respective crude birth rates (Table 1). The estimates are based on a prevalence for MFS of 2 – 3 per 10,000 individuals and the assumption that nucleotides in any location of the protein-coding region of *FBN1* are equally likely to be altered [16]. Given a US population of 331.5 M and 3.6 M newborns in 2020, we estimated a range from 82 – 123 newborns and 7,558 – 11,337 individuals with MFS due to a PTC variant in the US. The affected individuals worldwide could range from 3,210 – 4,815 newborns and 177,612 – 266,418 individuals. The worldwide estimates were based on a population of 7,790 M and a crude birth rate of 18.077/1,000 people. Collectively, this suggests that a substantial subpopulation of individuals with MFS could be considered for translational readthrough therapy.

**Table 1.** Estimated number of individuals with Marfan syndrome (MFS) due to a premature termination codon (PTC) in *FBN1*

|  | Demographics<br>(2020) | Individuals with MFS<br>(prevalence: 2 – 3 per 10,000) | Individuals with MFS<br>due to PTC in *FBN1*<br>(11.4% of *FBN1* variants) |
| --- | --- | --- | --- |
| US Population<sup>1</sup> | 331.5M | 66,300 – 99,450 | 7,558 – 11,337 |
| US Newborns<sup>2</sup> | 3.6M | 720 – 1,080 | 82 – 123 |
| World Population<sup>3</sup> | 7,790M | 1,558,000 – 2,337,000 | 177,612 – 266,418 |
| World Newborns<sup>4</sup> | 140.8M | 28,160 – 42,240 | 3,210 – 4,815 |

<sup>1</sup>Based on data from the US Census Bureau. <sup>2</sup>Based on data from the Center for Disease Control and Prevention (National Center for Health Statistics). <sup>3</sup>Based on data from the US Census Bureau. <sup>4</sup>Based on an estimated crude birth rate of 18.077 newborns/1000 people (Macrotrends.net).

#### Small molecules that block NMD, but not translational readthrough drugs, showed toxicity in MFS patient-derived fibroblasts

To experimentally test if targeting translational readthrough with or without suppression of NMD could ameliorate FBN1 deposition in the ECM, we obtained three previously characterized MFS patient-derived primary skin fibroblast lines from the Coriell Institute for Medical Research (Table 2) [30]. The cell lines represent different locations of the PTC in the *FBN1* sequence, different amino acid codons changed into PTCs and different nucleotide changes, resulting in different types of PTCs (Fig 1B, top). We first tested the effect of a single dose of small molecules that promote translational readthrough (ataluren, G418, gentamicin, amlexanox) or block NMD (amlexanox, NMDI 14) individually or in combination on all three MFS fibroblast lines (Table 3) [42–49].

**Table 2.** Characteristics of Marfan syndrome patient-derived primary skin fibroblasts^1^

| Coriell ID | PTC<br>Exon | *FBN1*<br>Variant<sup>1</sup> | Amino<br>acid<br>change<sup>1</sup> | PTC<br>(+1) | Age at sampling<br>in years<br>(Sex) | Mutant<br>transcript<br>level<sup>2</sup> | Group<sup>3</sup> |
| --- | --- | --- | --- | --- | --- | --- | --- |
| GM21966 | 7 | c.745G>T | p.E249* | <u>uaau</u> | 67 (male) | Marked<br>reduction | II |
| GM21970 | 15 | c.1884C>A | p.C628* | <u>ugag</u> | 57 (female) | Severe<br>reduction | I |
| GM21983 | 39 | c.4930C>T | p.R1644* | <u>ugag</u> | 38 (male) | n.d. | II |

<sup>1</sup>Nomenclature is based on the *FBN1* sequence NM\_000138.5, with the first nucleotide of the start codon being 1 and NP\_000129.3. <sup>2</sup>*FBN1* mutations and *FBN1* secretion and deposition phenotypes were extracted from Schrijver et al, 2002. <sup>3</sup>The biosynthesis groups described in Schrijver et al are based on Aoyama et al, 1994: Group I: Decreased *FBN1* synthesis but normal ECM deposition; Group II: Decreased *FBN1* synthesis and decreased ECM deposition.

**Table 3.** Small molecules used in this study

| Compound | Mode of action | Solvent | Source | References |
| --- | --- | --- | --- | --- |
| Amlexanox | Increases amount of PTC-containing mRNA<br>Promotes translational readthrough | DMSO | Sigma-Aldrich (SML0517) | Gonzalez-Hilarion et al, 2012<br>Atanasova et al, 2017 |
| Ataluren (PTC124) | Promotes translational read-through | DMSO | Selleckchem (S6003) | Wilschanski et al, 2011<br>Finkel, 2010 |
| G418 (Geneticin) | Inhibits elongation step during protein synthesis | PBS | Life Technologies (10131035) | Barton-Davis et al, 1999 |
| Gentamicin | Inhibits protein synthesis by binding to 30S ribosome subunit | water | Sigma-Aldrich (G1397) | Clancy et al, 2001 |
| NMDI 14 | Inhibits nonsense-mediated mRNA decay | DMSO | Sigma-Aldrich (SML1538) | Martin et al, 2014 |

DMSO: Dimethyl sulfoxide; PBS: Phosphate-buffered saline; PTC, premature termination codon

FBN1 and fibronectin deposition into the ECM after small molecule treatment was evaluated by immunofluorescence staining. Fibronectin was monitored since it forms a template that is required for FBN1 deposition in fibroblasts [50]. Ataluren and gentamicin were well tolerated and we observed a variable but consistent increase of FBN1 deposition with gentamicin in all three fibroblast cell lines and cell line dependent changes in FBN1 deposition with ataluren (Fig 2A, B and data not shown). In contrast, treatment of MFS fibroblasts with G418 reduced FBN1 and fibronectin deposition and resulted in apparent intracellular fibronectin retention. When the NMD inhibitor NMDI 14 was added to MFS fibroblasts alone or in combination with ataluren or G418, the cell number was greatly reduced and only sparse FBN1 and fibronectin deposition was observed (Fig 2C, D and data not shown). This suggested cytotoxicity of NMDI 14 in MFS fibroblasts at the concentration that we used. Treatment with amlexanox did not affect cell number, but fibronectin and concomitant FBN1 deposition was almost completely abolished compared to DMSO-treated controls for unclear reasons. Based on these data, we performed subsequent dose-response studies with ataluren and gentamicin.

**Fig 2.**
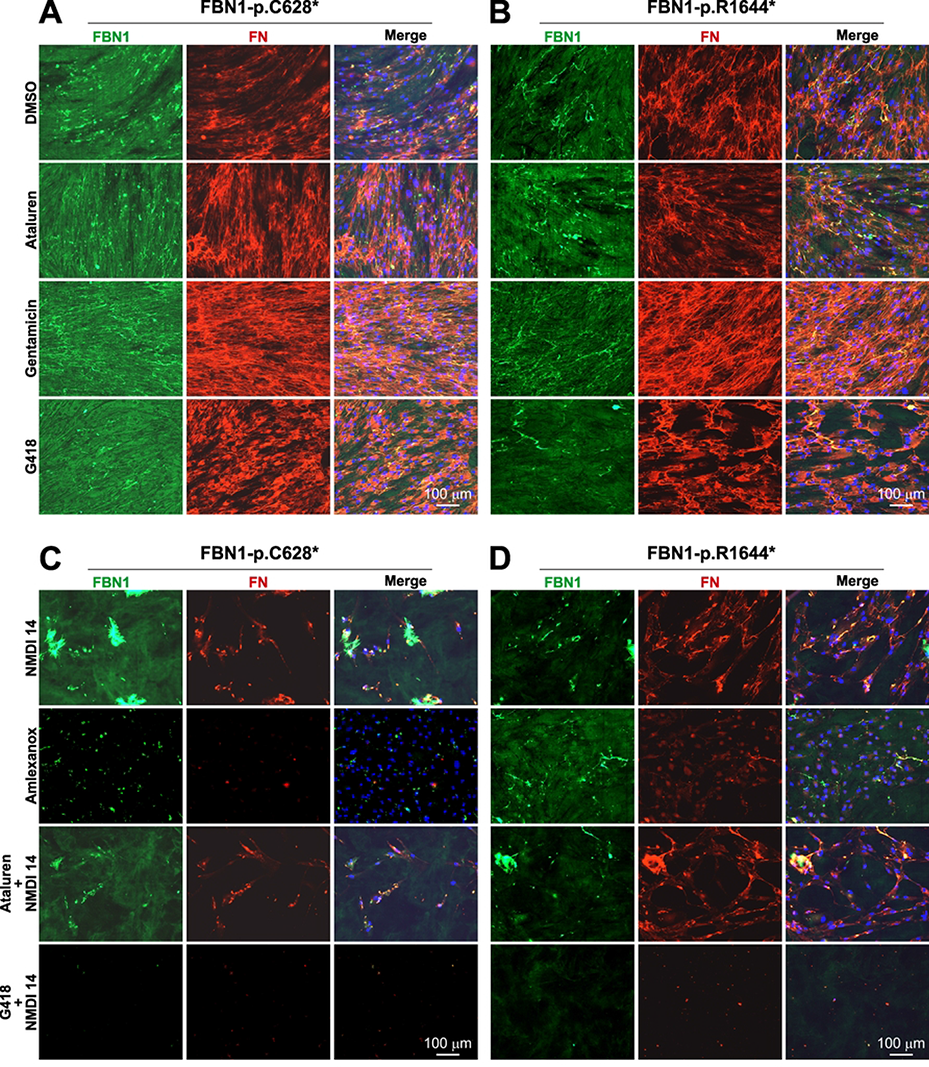
Treatment of MFS fibroblasts with small molecules promoting translational readthrough or inhibiting NMD. **(A, B)** Immunostaining for FBN1 and fibronectin deposition after treatment of fibroblasts harboring the FBN1-p.C628* (A) or FBN1-p.R1644* (B) mutation with small molecules promoting translational readthrough. Note reduced FBN1 deposition and aberrant fibronectin deposition with intracellular retention in the presence of G418; **(C, D)** FBN1 and fibronectin deposition after treatment of fibroblasts harboring the FBN1-p.C628* (C) or FBN1-p.R1644* (D) variant with NMD inhibitors individually or in combination with small molecules promoting translational readthrough. Representative images of n=2 experiments are shown.

#### Dose-dependent ataluren- or gentamicin-induced FBN1 deposition was cell-line specific

We determined the dose-dependent response of FBN1 and fibronectin deposition after treatment with increasing concentrations of ataluren (0 – 50 ⍰M) or gentamicin (0 – 2 mM) and observed variable responses of the three MFS fibroblast lines likely due to the specific FBN1 variant or donor-dependent effects. When fibroblasts harboring the FBN1-p.E249* variant were treated with ataluren, an increase in FBN1 and fibronectin deposition was observed with 5 ⍰M (Fig 3A-C). However, FBN1, but not FN deposition decreased at higher concentrations when normalized to the DAPI signal, and the deposition pattern became aberrant after treatment with 50 ⍰M ataluren. Western blot analysis suggested a reduction of FBN1 protein in conditioned medium and cell lysates, while fibronectin was affected to a lesser extent in the medium and no changes were detected in cell lysates (Fig 3D, E). This reduction may be attributed to increased deposition of FBN1 in the ECM or reduced extractability of the ECM from the cell lysate/ECM layer. After gentamicin treatment of fibroblasts harboring the FBN1-p.E249* variant, FBN1 deposition was increased in a dose-dependent manner, plateauing at 1 mM (Fig 3F-H). Fibronectin was increased and plateaued at 0.2 mM gentamicin. No changes in FBN1 or fibronectin protein in the medium or cell lysates were observed by western blot analysis (Fig 3I, J).

**Fig 3.**
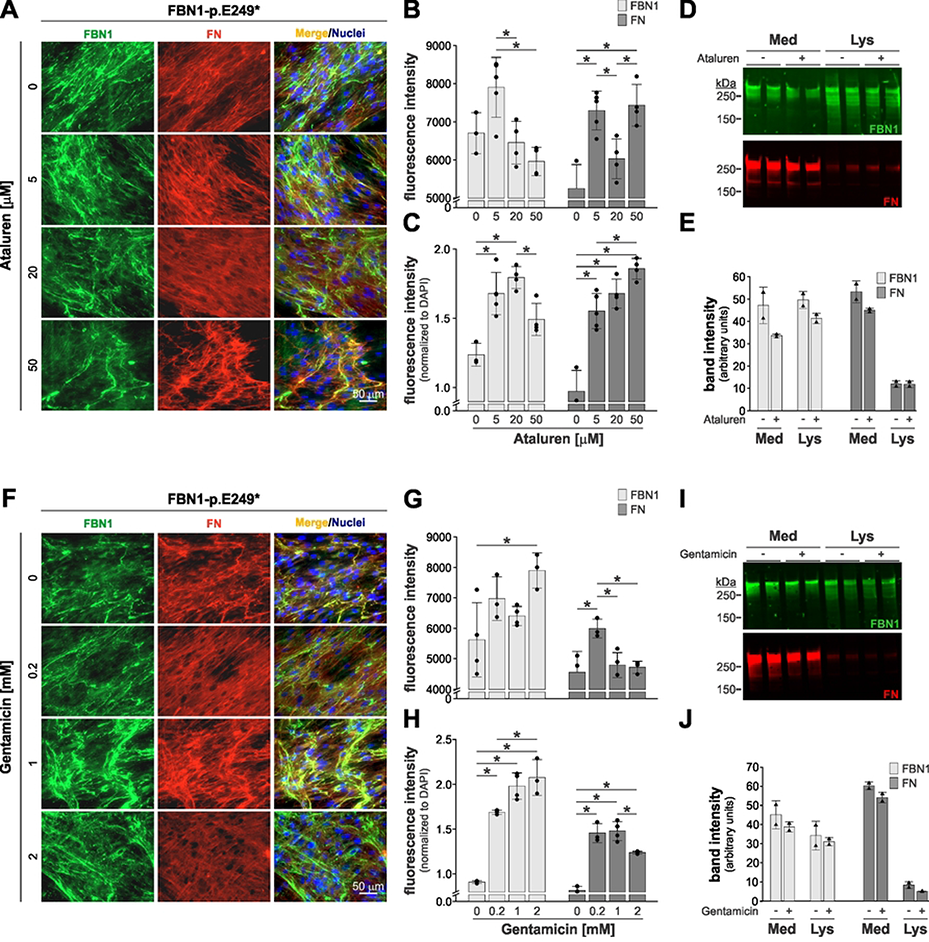
Dose-response studies with MFS fibroblasts harboring the FBN1-p.E249* variant. **(A)** Immunostaining for FBN1 and fibronectin deposition after treatment of fibroblasts harboring the FBN1-p.E249* variant with increasing concentrations of ataluren; **(B, C)** Quantification of fluorescence intensity of the FBN1 and fibronectin channel after ataluren treatment without (B) and with (C) normalization to DAPI signal; **(D)** Western blot analysis for FBN1 and fibronectin in conditioned medium (Med) and cell lysates (Lys) after treatment with ataluren; **(E)** Quantification of fluorescence intensity of the major FBN1 and fibronectin band; **(F)** Immunostaining for FBN1 and fibronectin deposition after treatment of fibroblasts harboring the FBN1-p.E249* variant with increasing concentrations of gentamicin; **(G, H)** Quantification of fluorescence intensity of the FBN1 and fibronectin channel after gentamicin treatment without (G) and with (H) normalization to DAPI signal. **(I)** Western blot analysis for FBN1 and fibronectin of conditioned medium (Med) and cell lysates (Lys) after treatment with gentamicin; **(J)** Quantification of fluorescence intensity of the major FBN1 and fibronectin band. Representative images from n=2 experiments are shown. 3-4 fields of view were quantified in panels B, C and G, H. Asterisks indicated p<0.05 based on a one-way ANOVA with posthoc Tukey test.

We also analyzed FBN1 and fibronectin deposition in fibroblasts harboring the FBN1-p.C628* or FBN1-p.R1644* variant after ataluren and gentamicin treatment by immunofluorescence staining. We found an increase in FBN1 fluorescence intensity when normalized to DAPI after FBN1-p.C628* fibroblasts were treated with 5 or 20 ⍰M ataluren, but not with gentamicin (Fig 4A-D). When fibroblasts harboring the FBN1-p.R1644* variant were treated with ataluren we observed no difference in FBN1 or FN fluorescence intensity (Fig 5A, B). When FBN1-p.R1644* fibroblasts were treated with gentamicin, we observed a dose-dependent reduction of the normalized FBN1 signal, but no change in FN deposition, suggesting a potential deleterious effect (Fig 5C, D).

**Fig 4.**
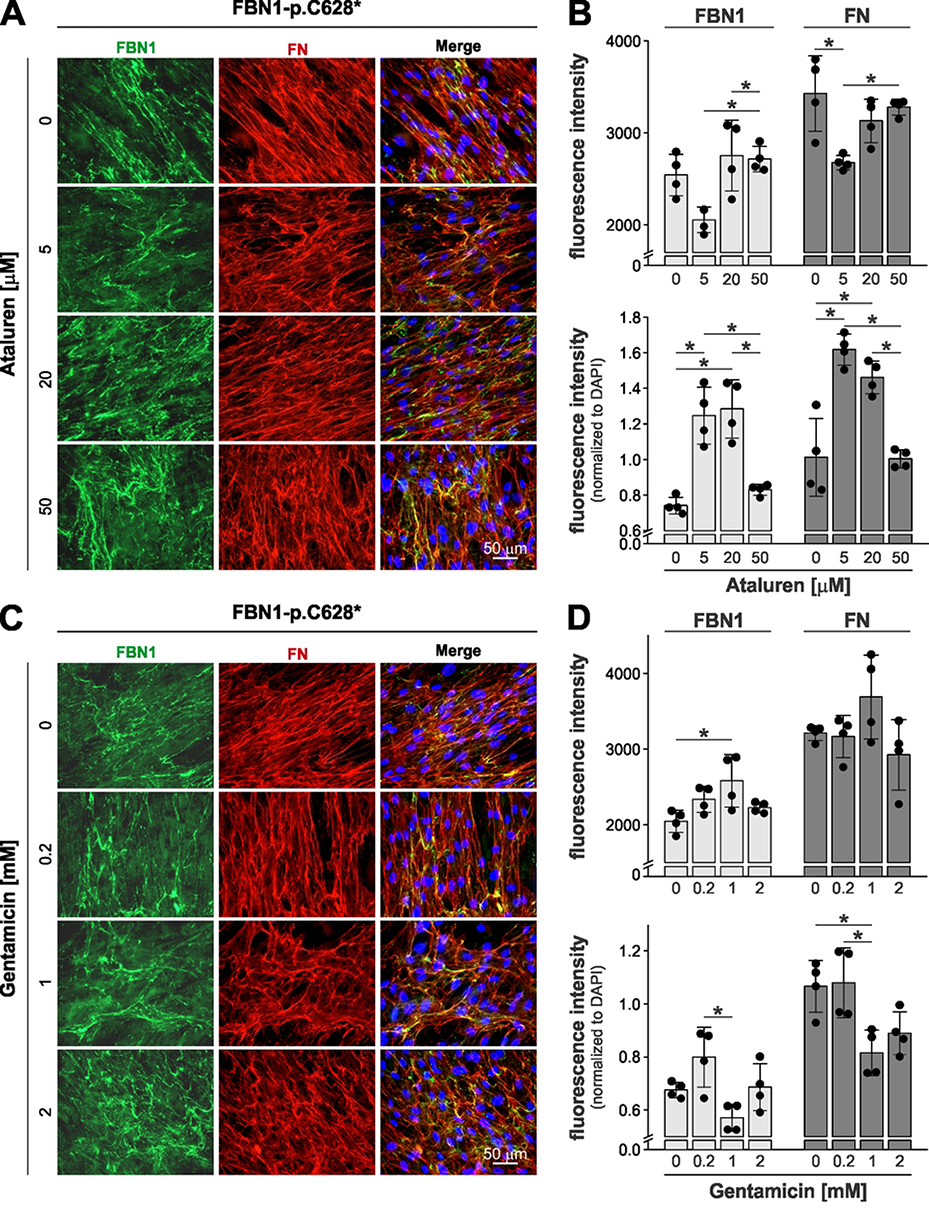
Dose-response studies with MFS fibroblasts harboring the FBN1-p.C628* variant. **(A)** Immunostaining for FBN1 and fibronectin deposition after treatment of fibroblasts harboring the FBN1-p.C628* variant with increasing concentrations of ataluren; **(B)** Quantification of fluorescence intensity of the FBN1 and fibronectin channel after ataluren treatment without (top) and with (bottom) normalization to DAPI signal; **(C)** Immunostaining for FBN1 and fibronectin deposition after treatment of fibroblasts harboring the FBN1-p.C628* variant with increasing concentrations of gentamicin; **(D)** Quantification of fluorescence intensity of the FBN1 and fibronectin channel after gentamicin treatment without (top) and with (bottom) normalization to cell number. Representative images from n=2 experiments are shown. 3-4 fields of view were quantified in panels B, D. Asterisks indicated p<0.05 based on a one-way ANOVA with posthoc Tukey test.

**Fig 5.**
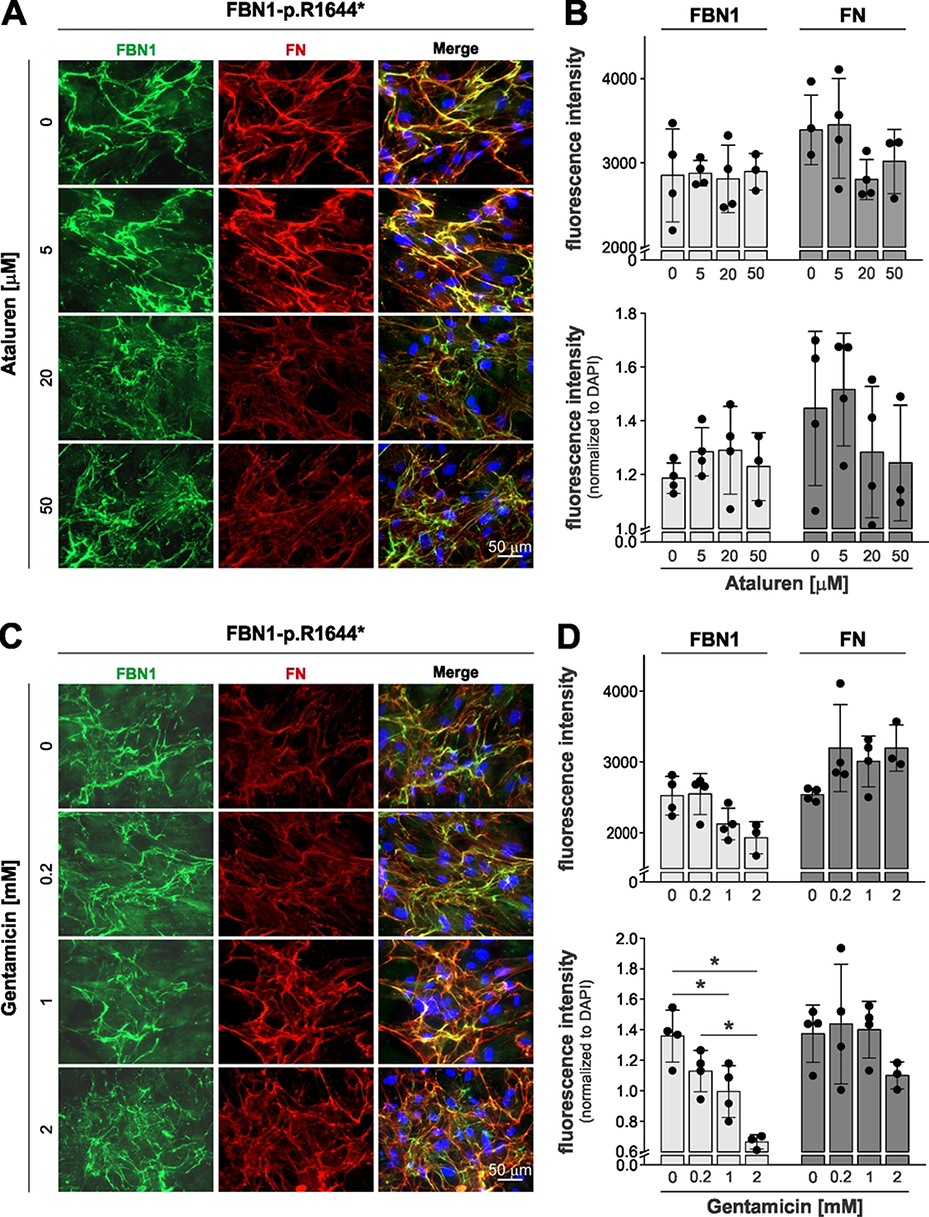
Dose-response studies with MFS fibroblasts harboring the FBN1-p.R1644* variant. **(A)** Immunostaining for FBN1 and fibronectin deposition after treatment of fibroblasts harboring the FBN1-p.R1644* variant with increasing concentrations of ataluren; **(B)** Quantification of fluorescence intensity of the FBN1 and fibronectin channel after ataluren treatment without (top) and with (bottom) normalization to DAPI signal; **(C)** Immunostaining for FBN1 and fibronectin deposition after treatment of fibroblasts harboring the FBN1-p.R1644* variant with increasing concentrations of gentamicin; **(D)** Quantification of fluorescence intensity of the FBN1 and fibronectin channel after gentamicin treatment without (top) and with (bottom) normalization to cell number. Representative images from n=2 experiments are shown. 3-4 fields of view were quantified in panels B, D. Asterisks indicated p<0.05 based on a one-way ANOVA with posthoc Tukey test.

Collectively, these data suggest that promoting translational readthrough in MFS fibroblasts resulted in a likely PTC and drug-specific response where ECM deposition of FBN1 was increased for some PTC variants.

## Discussion

Due to the large number of pathogenic variants in *FBN1* an opportunity may exist to design personalized treatment approaches that may be beneficial for a subset of individuals with MFS due to specific *FBN1* variants, such as PTCs, which account for 11.4% of all MFS-causing *FBN1* variants. Here, we provide some evidence that suppression of PTCs with small molecules that promote translational readthrough may result in enhanced deposition of FBN1 in the ECM of MFS patient-derived fibroblasts. However, this effect was cell line dependent and its magnitude was likely determined by the specific PTC variant in *FBN1*. In addition, factors such as age of the patient at the time of sampling, which ranged from 22 to 67 years of age, and the individual genetic background may also be sources of the observed variability. This highlights the likely need for a personalized medicine approach with preclinical testing of individual variants in patient-derived cells or organoids to determine if translational readthrough therapy can be considered for individual MFS patients.

The correlation of specific *FBN1* variants with the patient-specific expression of the MFS phenotype remains a key challenge [24, 38, 51–53]. However, it has been previously shown that PTC variants in *FBN1* resulted in exacerbated skeletal features but a reduced risk for ocular involvement [30]. In addition, dissection of the ascending aorta was more prevalent in individuals with PTC variants in *FBN1*. A more recent study confirmed these findings and demonstrated that a cohort of individuals with MFS due to haploinsufficient *FBN1* variants, including PTCs, showed a more rapid rate of aortic dilatation [54]. As such, PTC variants in *FBN1* may be promoting a more rapid progression of aortic dilatation, which is the most critical symptom to monitor in order to prevent aortic dissection or rupture by preemptively replacing the dilated part of the aorta with a vascular graft [55]. Therefore, strategies to suppress PTCs and augment FBN1 synthesis by promoting translational readthrough may be a therapeutic approach to be considered and developed further.

Translational readthrough therapy is currently in preclinical or clinical trials for several genetic disorders, such as cystic fibrosis, Duchenne muscular dystrophy or epidermolysis bullosa [56–59]. The premise for this approach is that by promoting translational readthrough through PTCs, a threshold amount of functional protein, sufficient to ameliorate disease outcome, can be generated. However, several factors influence the possible success of such a therapeutic approach. For example, most of the preclinical and clinical studies of translational readthrough therapy targeted recessive disorders, which presumably are caused by a severe reduction or complete absence of functional protein. However, MFS is an autosomal dominant disorder in which one wild-type *FBN1* allele is retained. As a consequence, the amount of FBN1 in the ECM is reduced by 50% or more depending on the nature of the individual *FBN1* variant being categorized as haploinsufficient or dominant-negative. However, it is unclear how much additional FBN1 protein would need to be deposited in the ECM to result in a therapeutic benefit for patients with MFS due to a PTC. In previous studies, augmentation of 3-10% of the original protein amount could be achieved by promoting translational readthrough in several unrelated genetic disorders [8]. However, most of these studies were performed without inhibition of NMD, which may be the key factor necessary to increase the starting amount of variant mRNA. Based on the categorization of MFS variants in five biochemical groups, PTC variants that showed reduced secretion but normal deposition would probably be the most receptive to promoting translational readthrough in order to augment FBN1 deposition [25]. In these cases, an increase in FBN1 protein is expected to translate into increased FBN1 deposition. In contrast, *FBN1* variants that compromise synthesis and ECM deposition may not respond to a further increase in FBN1 protein amount by enhancing FBN1 ECM deposition and in fact may result in exacerbated reduction of FBN1 deposition through a dominant-negative effect. Experimentally, the consequences of promoting translational readthrough on FBN1 deposition in the context of individual *FBN1* PTCs could be tested in cell culture using MFS patient-derived fibroblasts or induced pluripotent stem cells (iPSCs) differentiated into vascular smooth muscle cells or skin fibroblasts [60, 61].

As mentioned above, the success of a translational readthrough approach for MFS is likely dependent on the amount of *FBN1* mRNA harboring the PTC that would be available for translation. Unfortunately, the NMD inhibitors that we tested, amlexanox and NMDI 14, resulted in the absence of ECM deposition or toxicity towards the MFS fibroblasts, respectively. Therefore, we were not able to test a combination of NMD inhibition and PTC suppression on the effect of FBN1 deposition in the MFS patient-derived fibroblasts. Since amlexanox is a pleiotropic drug with multiple effects, it is unclear what caused the absence of ECM deposition, even though the cell number appeared not to be reduced [62]. Phenotypic variability even within families with the same *FBN1* variant is a known feature of MFS and complicates genotype-phenotype correlations. One explanation for this variability is the variability in the expression of the wild-type *FBN1* allele [63, 64]. An interesting alternative to inhibiting NMD would be to enhance the expression of the wild-type *FBN1* allele, specifically in the context of a PTC variant that would not exert a dominant-negative effect due to NMD of the variant allele. Such an approach would be conceptually feasible with the recent development of gene-specific CRISPR-based activation systems [65, 66]. If NMD of the variant *FBN1* allele can be inhibited, the success of a translational readthrough approach will further depend on the reinserted amino acid, which is determined by the individual PTC where UGA results in tyrosine, arginine or cysteine incorporation and UAA or UGA in glutamine, tyrosine or lysine incorporation [7, 15]. However, the ratio of the insertion of the three amino acids can be influenced by the experimental system and the choice of readthrough-inducing agent. This could be relevant to further stratify the population of MFS patients with PTCs and test different translational readthrough compounds to avoid a potentially dominant-negative effect of variant FBN1 due to the insertion of a deleterious amino acid in the PTC position. Such an effect could potentially result in the further reduction of FBN1 deposition and possibly in a worsening of the MFS phenotype or in accelerated disease progression. In our experiments, the FBN1-p.R1644* fibroblasts showed such a behavior and the amount of FBN1 that was deposited into the ECM was reduced with increasing doses of gentamicin. This particular variant resulted from a UGA PTC and the inclusion of a cysteine or tyrosine, in addition to the wild type arginine, which may interfere with FBN1 ECM deposition. However, a proteomics approach would be needed to identify the ratio of amino acids incorporated in FBN1-p.R1644* and different readthrough drugs would need to be tested to determine if this ratio can be changed in a way that would promote FBN1 deposition in the ECM and suppress a potential dominant-negative effect [15, 44].

Finally, it is worth considering under which condition a translational readthrough therapy would be the most beneficial. With the exception of the neonatal form, MFS is relatively slow progressing and thus offers a substantial therapeutic window where translational readthrough therapy could result in improved patient outcome. Most relevant would be slowing or arresting the growth of aortic aneurysms to delay or prevent the need for aortic surgery, reducing the risk for lens dislocation, and ameliorating musculoskeletal symptoms, which are the main sources of pain and mobility restrictions in patients with MFS. While considerable amounts of FBN1 are deposited in the ECM already at birth, further FBN1 deposition is likely required for postnatal growth and tissue homeostasis, despite the limited availability of experimental data [67]. Therefore, promoting FBN1 deposition during the growth phase may be beneficial. With the increased availability of affordable whole exome sequencing, potentially as part of routine newborn screenings in the future, many MFS patients may be identified before the onset of symptoms and thus the therapeutic window for translational readthrough therapy may be extended to the benefit of eligible MFS patients and allow for preclinical testing of individual MFS patients.

## Conclusions

Collectively, we demonstrate that promoting translational readthrough could be considered as a therapeutic approach for MFS for patients with specific PTCs in *FBN1*. Several limitations of our study would need to be addressed in future research: First, we analyzed the response to translational readthrough drugs of only four out of 210 PTCs that result in MFS and it is possible that each PTC in *FBN1* responds differently due to the position of the PTC within the exon, its sequence context and the residual amount of variant *FBN1* mRNA [41]. Second, we tested only three translational readthrough drugs, one of which (G418) did result in aberrant FBN1 deposition and intracellular fibronectin retention and there is some concern about the efficacy of ataluren [68]. Ataluren itself shows a bell shaped dose-response curve, where its efficacy decreases again at higher concentrations and the dosing for individual *FBN1* variants may need to be optimized [42, 69]. It is likely that the efficiency of individual drugs to promote translational readthrough of *FBN1* PTCs will vary depending on the specific PTC and the amino acids that are reinserted. However, it is anticipated that more small molecules will be developed as translational readthrough drugs in the future that could then be tested on MFS patient-derived fibroblasts. Third, the translational readthrough drugs that we used could have non-FBN1 effects on cell physiology and protein secretion that could confound FBN1 ECM deposition. Fourth, due to toxicity of the NMD inhibitors we were not able to test if blocking NMD in combination with promoting translational readthrough would synergistically augment FBN1 deposition in the ECM. It is important to note that systemically inhibiting NMD, which is an important endogenous quality control mechanism to prevent the accumulation of toxic protein products due to randomly occurring PTCs, may have unwanted side effects and may not be feasible in vivo. However, innovative antisense oligonucleotide technology was recently developed for gene-specific NMD inhibition that could be combine with readthrough therapy [70]. Finally, there is currently no mouse model for MFS due to a *FBN1* PTC available, where a translational readthrough approach for MFS could have been tested in vivo. Despite these limitations, we suggest that suppression of PTCs could be an innovative therapeutic approach for specific patients with MFS due to PTCs in *FBN1* (Fig. 6). By using defined metrics, such as FBN1 synthesis and deposition in MFS patient-derived fibroblasts exposed to translational readthrough drugs, a clinical decision to consider individual MFS patients for translational readthrough therapy could be made in a bedside-to-bench-to-bedside personalized medicine approach.

**Fig 6.**
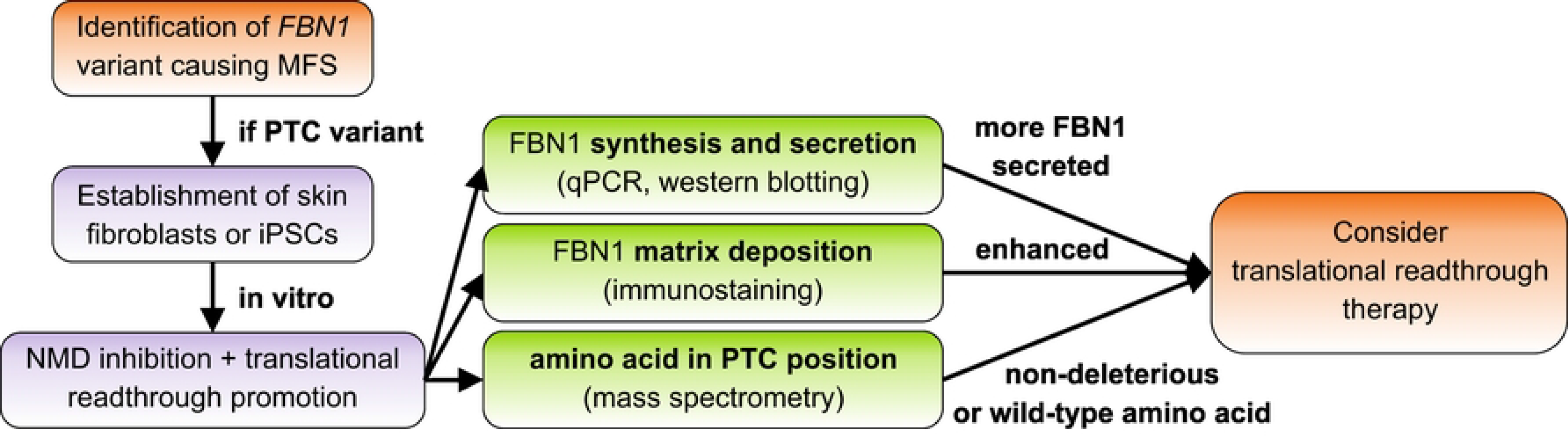
Possible decision tree to identify individuals with MFS that may benefit from PTC suppression. The decision tree is based on the identification of the PTC variant in *FBN1* in a patient with suspected or confirmed MFS and the identification of a beneficial effect of inhibiting NMD or promoting translational readthrough in patient-derived skin fibroblasts or induced pluripotent stem cells (iPSCs). Key parameters to be evaluated are increases in *FBN1* mRNA resulting in increased FBN1 protein secretion, increased FBN1 deposition in the ECM, and the identification of the amino acid that is incorporated in the position of the PTC. A clinical decision could even include the investigation of the individual PTC variant in iPSC-derived organoids or patient-specific *Fbn1* knock-in mouse models.

## Acknowledgments

We thank Dr. Dieter Reinhardt (McGill University, Montreal, Canada) for generously providing the FBN1 antibody.

## Author Contributions

Conceptualization, D.H; methodology, Z.B. and D.H.; formal analysis, Z.B. and D.H.; investigation, Z.B. and D.H.; writing—original draft preparation, D.H.; writing—review and editing, Z.B. and D.H.; visualization, D.H.; supervision, D.H.; funding acquisition, D.H. All authors have read and agreed to the published version of the manuscript.”

## Data Availability

Data underlying Fig. 1 were derived from the following resources available in the public domain: UMD *FBN1* database (http://www.umd.be/FBN1/) [19]. Data for Table 2 were extracted from [30].

## References

1. Mort M, Ivanov D, Cooper DN, Chuzhanova NA. A meta-analysis of nonsense mutations causing human genetic disease. Hum Mutat. 2008;29(8):1037–47. Epub 2008/05/06. doi: 10.1002/humu.20763. PubMed PMID: 18454449.

2. Popp MW, Maquat LE. Leveraging Rules of Nonsense-Mediated mRNA Decay for Genome Engineering and Personalized Medicine. Cell. 2016;165(6):1319–22. Epub 2016/06/04. doi: 10.1016/j.cell.2016.05.053. PubMed PMID: 27259145; PubMed Central PMCID: PMCPMC4924582.

3. Maquat LE, Kinniburgh AJ, Rachmilewitz EA, Ross J. Unstable beta-globin mRNA in mRNA-deficient beta o thalassemia. Cell. 1981;27(3 Pt 2):543–53. Epub 1981/12/01. doi: 10.1016/0092-8674(81)90396-2. PubMed PMID: 6101206.

4. Isken O, Maquat LE. Quality control of eukaryotic mRNA: safeguarding cells from abnormal mRNA function. Genes Dev. 2007;21(15):1833–56. Epub 2007/08/03. doi: 10.1101/gad.1566807. PubMed PMID: 17671086.

5. Malik V, Rodino-Klapac LR, Viollet L, Wall C, King W, Al-Dahhak R, et al. Gentamicin-induced readthrough of stop codons in Duchenne muscular dystrophy. Ann Neurol. 2010;67(6):771–80. Epub 2010/06/03. doi: 10.1002/ana.22024. PubMed PMID: 20517938.

6. Borgatti M, Altamura E, Salvatori F, D’Aversa E, Altamura N. Screening Readthrough Compounds to Suppress Nonsense Mutations: Possible Application to beta-Thalassemia. J Clin Med. 2020;9(2). Epub 2020/01/25. doi: 10.3390/jcm9020289. PubMed PMID: 31972957; PubMed Central PMCID: PMCPMC7073686.

7. Roy B, Leszyk JD, Mangus DA, Jacobson A. Nonsense suppression by near-cognate tRNAs employs alternative base pairing at codon positions 1 and 3. Proc Natl Acad Sci U S A. 2015;112(10):3038–43. Epub 2015/03/04. doi: 10.1073/pnas.1424127112. PubMed PMID: 25733896; PubMed Central PMCID: PMCPMC4364220.

8. Lombardi S, Testa MF, Pinotti M, Branchini A. Molecular Insights into Determinants of Translational Readthrough and Implications for Nonsense Suppression Approaches. Int J Mol Sci. 2020;21(24). Epub 2020/12/17. doi: 10.3390/ijms21249449. PubMed PMID: 33322589; PubMed Central PMCID: PMCPMC7764779.

9. Martins-Dias P, Romao L. Nonsense suppression therapies in human genetic diseases. Cell Mol Life Sci. 2021;78(10):4677–701. Epub 2021/03/23. doi: 10.1007/s00018-021-03809-7. PubMed PMID: 33751142.

10. Burke JF, Mogg AE. Suppression of a nonsense mutation in mammalian cells in vivo by the aminoglycoside antibiotics G-418 and paromomycin. Nucleic Acids Res. 1985;13(17):6265–72. Epub 1985/09/11. doi: 10.1093/nar/13.17.6265. PubMed PMID: 2995924; PubMed Central PMCID: PMCPMC321951.

11. Wilschanski M, Miller LL, Shoseyov D, Blau H, Rivlin J, Aviram M, et al. Chronic ataluren (PTC124) treatment of nonsense mutation cystic fibrosis. Eur Respir J. 2011;38(1):59–69. Epub 2011/01/15. doi: 10.1183/09031936.00120910. PubMed PMID: 21233271.

12. Finkel RS. Read-through strategies for suppression of nonsense mutations in Duchenne/ Becker muscular dystrophy: aminoglycosides and ataluren (PTC124). J Child Neurol. 2010;25(9):1158–64. Epub 2010/06/04. doi: 10.1177/0883073810371129. PubMed PMID: 20519671; PubMed Central PMCID: PMCPMC3674569.

13. Gonzalez-Hilarion S, Beghyn T, Jia J, Debreuck N, Berte G, Mamchaoui K, et al. Rescue of nonsense mutations by amlexanox in human cells. Orphanet J Rare Dis. 2012;7:58. Epub 2012/09/04. doi: 10.1186/1750-1172-7-58. PubMed PMID: 22938201; PubMed Central PMCID: PMCPMC3562214.

14. Pranke I, Bidou L, Martin N, Blanchet S, Hatton A, Karri S, et al. Factors influencing readthrough therapy for frequent cystic fibrosis premature termination codons. ERJ Open Res. 2018;4(1). Epub 2018/03/03. doi: 10.1183/23120541.00080-2017. PubMed PMID: 29497617; PubMed Central PMCID: PMCPMC5827411.

15. Blanchet S, Cornu D, Argentini M, Namy O. New insights into the incorporation of natural suppressor tRNAs at stop codons in Saccharomyces cerevisiae. Nucleic Acids Res. 2014;42(15):10061–72. Epub 2014/07/25. doi: 10.1093/nar/gku663. PubMed PMID: 25056309; PubMed Central PMCID: PMCPMC4150775.

16. Loeys BL, Dietz HC, Braverman AC, Callewaert BL, De Backer J, Devereux RB, et al. The revised Ghent nosology for the Marfan syndrome. J Med Genet. 2010;47(7):476–85. Epub 2010/07/02. doi: 10.1136/jmg.2009.072785. PubMed PMID: 20591885.

17. Pyeritz RE. Marfan syndrome: improved clinical history results in expanded natural history. Genet Med. 2019;21(8):1683–90. Epub 2018/12/24. doi: 10.1038/s41436-018-0399-4. PubMed PMID: 30573797.

18. Milewicz DM, Prakash SK, Ramirez F. Therapeutics Targeting Drivers of Thoracic Aortic Aneurysms and Acute Aortic Dissections: Insights from Predisposing Genes and Mouse Models. Annu Rev Med. 2017;68:51–67. Epub 2017/01/19. doi: 10.1146/annurev-med-100415-022956. PubMed PMID: 28099082; PubMed Central PMCID: PMCPMC5499376.

19. Collod-Beroud G, Le Bourdelles S, Ades L, Ala-Kokko L, Booms P, Boxer M, et al. Update of the UMD-FBN1 mutation database and creation of an FBN1 polymorphism database. Hum Mutat. 2003;22(3):199–208. Epub 2003/08/26. doi: 10.1002/humu.10249. PubMed PMID: 12938084.

20. Ramirez F, Rifkin DB. Extracellular microfibrils: contextual platforms for TGFbeta and BMP signaling. Curr Opin Cell Biol. 2009;21(5):616–22. Epub 2009/06/16. doi: 10.1016/j.ceb.2009.05.005. PubMed PMID: 19525102; PubMed Central PMCID: PMCPMC2767232.

21. Zimmermann LA, Correns A, Furlan AG, Spanou CES, Sengle G. Controlling BMP growth factor bioavailability: The extracellular matrix as multi skilled platform. Cell Signal. 2021;85:110071. Epub 2021/07/05. doi: 10.1016/j.cellsig.2021.110071. PubMed PMID: 34217834.

22. Thomson J, Singh M, Eckersley A, Cain SA, Sherratt MJ, Baldock C. Fibrillin microfibrils and elastic fibre proteins: Functional interactions and extracellular regulation of growth factors. Seminars in cell & developmental biology. 2019;89:109–17. Epub 2018/07/18. doi: 10.1016/j.semcdb.2018.07.016. PubMed PMID: 30016650; PubMed Central PMCID: PMCPMC6461133.

23. Milewicz DM, Ramirez F. Therapies for Thoracic Aortic Aneurysms and Acute Aortic Dissections. Arterioscler Thromb Vasc Biol. 2019;39(2):126–36. Epub 2019/01/18. doi: 10.1161/ATVBAHA.118.310956. PubMed PMID: 30651002; PubMed Central PMCID: PMCPMC6398943.

24. Faivre L, Collod-Beroud G, Loeys BL, Child A, Binquet C, Gautier E, et al. Effect of mutation type and location on clinical outcome in 1,013 probands with Marfan syndrome or related phenotypes and FBN1 mutations: an international study. Am J Hum Genet. 2007;81(3):454–66. Epub 2007/08/19. doi: 10.1086/520125. PubMed PMID: 17701892; PubMed Central PMCID: PMCPMC1950837.

25. Aoyama T, Francke U, Dietz HC, Furthmayr H. Quantitative differences in biosynthesis and extracellular deposition of fibrillin in cultured fibroblasts distinguish five groups of Marfan syndrome patients and suggest distinct pathogenetic mechanisms. J Clin Invest. 1994;94(1):130–7. Epub 1994/07/01. doi: 10.1172/JCI117298. PubMed PMID: 8040255; PubMed Central PMCID: PMCPMC296290.

26. Judge DP, Biery NJ, Keene DR, Geubtner J, Myers L, Huso DL, et al. Evidence for a critical contribution of haploinsufficiency in the complex pathogenesis of Marfan syndrome. J Clin Invest. 2004;114(2):172–81. Epub 2004/07/16. doi: 10.1172/JCI20641. PubMed PMID: 15254584; PubMed Central PMCID: PMCPMC449744.

27. Zeyer KA, Reinhardt DP. Engineered mutations in fibrillin-1 leading to Marfan syndrome act at the protein, cellular and organismal levels. Mutat Res Rev Mutat Res. 2015;765:7–18. Epub 2015/08/19. doi: 10.1016/j.mrrev.2015.04.002. PubMed PMID: 26281765.

28. Potter KA, Hoffman Y, Sakai LY, Byers PH, Besser TE, Milewicz DM. Abnormal fibrillin metabolism in bovine Marfan syndrome. Am J Pathol. 1993;142(3):803–10. Epub 1993/03/01. PubMed PMID: 8456941; PubMed Central PMCID: PMCPMC1886805.

29. Milewicz DM, Pyeritz RE, Crawford ES, Byers PH. Marfan syndrome: defective synthesis, secretion, and extracellular matrix formation of fibrillin by cultured dermal fibroblasts. J Clin Invest. 1992;89(1):79–86. Epub 1992/01/01. doi: 10.1172/JCI115589. PubMed PMID: 1729284; PubMed Central PMCID: PMCPMC442822.

30. Schrijver I, Liu W, Odom R, Brenn T, Oefner P, Furthmayr H, et al. Premature termination mutations in FBN1: distinct effects on differential allelic expression and on protein and clinical phenotypes. Am J Hum Genet. 2002;71(2):223–37. Epub 2002/06/18. doi: 10.1086/341581. PubMed PMID: 12068374; PubMed Central PMCID: PMCPMC379156.

31. Tiedemann K, Batge B, Muller PK, Reinhardt DP. Interactions of fibrillin-1 with heparin/heparan sulfate, implications for microfibrillar assembly. J Biol Chem. 2001;276(38):36035–42. Epub 2001/07/20. doi: 10.1074/jbc.M104985200. PubMed PMID: 11461921.

32. Faivre L, Collod-Beroud G, Callewaert B, Child A, Binquet C, Gautier E, et al. Clinical and mutation-type analysis from an international series of 198 probands with a pathogenic FBN1 exons 24-32 mutation. Eur J Hum Genet. 2009;17(4):491–501. Epub 2008/11/13. doi: 10.1038/ejhg.2008.207. PubMed PMID: 19002209; PubMed Central PMCID: PMCPMC2734964.

33. Romere C, Duerrschmid C, Bournat J, Constable P, Jain M, Xia F, et al. Asprosin, a Fasting-Induced Glucogenic Protein Hormone. Cell. 2016;165(3):566–79. Epub 2016/04/19. doi: 10.1016/j.cell.2016.02.063. PubMed PMID: 27087445; PubMed Central PMCID: PMCPMC4852710.

34. Passarge E, Robinson PN, Graul-Neumann LM. Marfanoid-progeroid-lipodystrophy syndrome: a newly recognized fibrillinopathy. Eur J Hum Genet. 2016;24(9):1244–7. Epub 2016/02/11. doi: 10.1038/ejhg.2016.6. PubMed PMID: 26860060; PubMed Central PMCID: PMCPMC4989216.

35. Stanley S, Balic Z, Hubmacher D. Acromelic dysplasias: how rare musculoskeletal disorders reveal biological functions of extracellular matrix proteins. Ann N Y Acad Sci. 2020. Epub 2020/09/04. doi: 10.1111/nyas.14465. PubMed PMID: 32880985.

36. Le Goff C, Mahaut C, Wang LW, Allali S, Abhyankar A, Jensen S, et al. Mutations in the TGFbeta binding-protein-like domain 5 of FBN1 are responsible for acromicric and geleophysic dysplasias. Am J Hum Genet. 2011;89(1):7–14. Epub 2011/06/21. doi: 10.1016/j.ajhg.2011.05.012. PubMed PMID: 21683322; PubMed Central PMCID: PMCPMC3135800.

37. Nagy E, Maquat LE. A rule for termination-codon position within intron-containing genes: when nonsense affects RNA abundance. Trends Biochem Sci. 1998;23(6):198–9. Epub 1998/06/30. doi: 10.1016/s0968-0004(98)01208-0. PubMed PMID: 9644970.

38. Tiecke F, Katzke S, Booms P, Robinson PN, Neumann L, Godfrey M, et al. Classic, atypically severe and neonatal Marfan syndrome: twelve mutations and genotype-phenotype correlations in FBN1 exons 24-40. Eur J Hum Genet. 2001;9(1):13–21. Epub 2001/02/15. doi: 10.1038/sj.ejhg.5200582. PubMed PMID: 11175294.

39. Gerber EE, Gallo EM, Fontana SC, Davis EC, Wigley FM, Huso DL, et al. Integrin-modulating therapy prevents fibrosis and autoimmunity in mouse models of scleroderma. Nature. 2013;503(7474):126–30. Epub 2013/10/11. doi: 10.1038/nature12614. PubMed PMID: 24107997; PubMed Central PMCID: PMCPMC3992987.

40. Manuvakhova M, Keeling K, Bedwell DM. Aminoglycoside antibiotics mediate context-dependent suppression of termination codons in a mammalian translation system. RNA. 2000;6(7):1044–55. Epub 2000/08/05. doi: 10.1017/s1355838200000716. PubMed PMID: 10917599; PubMed Central PMCID: PMCPMC1369979.

41. Bidou L, Hatin I, Perez N, Allamand V, Panthier JJ, Rousset JP. Premature stop codons involved in muscular dystrophies show a broad spectrum of readthrough efficiencies in response to gentamicin treatment. Gene Ther. 2004;11(7):619–27. Epub 2004/02/20. doi: 10.1038/sj.gt.3302211. PubMed PMID: 14973546.

42. Welch EM, Barton ER, Zhuo J, Tomizawa Y, Friesen WJ, Trifillis P, et al. PTC124 targets genetic disorders caused by nonsense mutations. Nature. 2007;447(7140):87–91. Epub 2007/04/24. doi: 10.1038/nature05756. PubMed PMID: 17450125.

43. Roy B, Friesen WJ, Tomizawa Y, Leszyk JD, Zhuo J, Johnson B, et al. Ataluren stimulates ribosomal selection of near-cognate tRNAs to promote nonsense suppression. Proc Natl Acad Sci U S A. 2016;113(44):12508–13. Epub 2016/11/03. doi: 10.1073/pnas.1605336113. PubMed PMID: 27702906; PubMed Central PMCID: PMCPMC5098639.

44. Ng MY, Li H, Ghelfi MD, Goldman YE, Cooperman BS. Ataluren and aminoglycosides stimulate read-through of nonsense codons by orthogonal mechanisms. Proc Natl Acad Sci U S A. 2021;118(2). Epub 2021/01/09. doi: 10.1073/pnas.2020599118. PubMed PMID: 33414181; PubMed Central PMCID: PMCPMC7812769.

45. Barton-Davis ER, Cordier L, Shoturma DI, Leland SE, Sweeney HL. Aminoglycoside antibiotics restore dystrophin function to skeletal muscles of mdx mice. J Clin Invest. 1999;104(4):375–81. Epub 1999/08/17. doi: 10.1172/JCI7866. PubMed PMID: 10449429; PubMed Central PMCID: PMCPMC481050.

46. Clancy JP, Bebok Z, Ruiz F, King C, Jones J, Walker L, et al. Evidence that systemic gentamicin suppresses premature stop mutations in patients with cystic fibrosis. Am J Respir Crit Care Med. 2001;163(7):1683–92. Epub 2001/06/13. doi: 10.1164/ajrccm.163.7.2004001. PubMed PMID: 11401894.

47. Martin L, Grigoryan A, Wang D, Wang J, Breda L, Rivella S, et al. Identification and characterization of small molecules that inhibit nonsense-mediated RNA decay and suppress nonsense p53 mutations. Cancer Res. 2014;74(11):3104–13. Epub 2014/03/26. doi: 10.1158/0008-5472.CAN-13-2235. PubMed PMID: 24662918; PubMed Central PMCID: PMCPMC4040335.

48. Atanasova VS, Jiang Q, Prisco M, Gruber C, Pinon Hofbauer J, Chen M, et al. Amlexanox Enhances Premature Termination Codon Read-Through in COL7A1 and Expression of Full Length Type VII Collagen: Potential Therapy for Recessive Dystrophic Epidermolysis Bullosa. J Invest Dermatol. 2017;137(9):1842–9. Epub 2017/05/28. doi: 10.1016/j.jid.2017.05.011. PubMed PMID: 28549954; PubMed Central PMCID: PMCPMC5573171.

49. Reilly SM, Chiang SH, Decker SJ, Chang L, Uhm M, Larsen MJ, et al. An inhibitor of the protein kinases TBK1 and IKK-varepsilon improves obesity-related metabolic dysfunctions in mice. Nat Med. 2013;19(3):313–21. Epub 2013/02/12. doi: 10.1038/nm.3082. PubMed PMID: 23396211; PubMed Central PMCID: PMCPMC3594079.

50. Sabatier L, Chen D, Fagotto-Kaufmann C, Hubmacher D, McKee MD, Annis DS, et al. Fibrillin assembly requires fibronectin. Mol Biol Cell. 2009;20(3):846–58. Epub 2008/11/28. doi: 10.1091/mbc.E08-08-0830. PubMed PMID: 19037100; PubMed Central PMCID: PMCPMC2633374.

51. Stark VC, Hensen F, Kutsche K, Kortum F, Olfe J, Wiegand P, et al. Genotype-Phenotype Correlation in Children: The Impact of FBN1 Variants on Pediatric Marfan Care. Genes (Basel). 2020;11(7). Epub 2020/07/19. doi: 10.3390/genes11070799. PubMed PMID: 32679894; PubMed Central PMCID: PMCPMC7397236.

52. Groth KA, Von Kodolitsch Y, Kutsche K, Gaustadnes M, Thorsen K, Andersen NH, et al. Evaluating the quality of Marfan genotype-phenotype correlations in existing FBN1 databases. Genet Med. 2017;19(7):772–7. Epub 2016/12/03. doi: 10.1038/gim.2016.181. PubMed PMID: 27906200.

53. Hernandiz A, Zuniga A, Valera F, Domingo D, Ontoria-Oviedo I, Mari JF, et al. Genotype FBN1/phenotype relationship in a cohort of patients with Marfan syndrome. Clin Genet. 2021;99(2):269–80. Epub 2020/11/12. doi: 10.1111/cge.13879. PubMed PMID: 33174221.

54. Franken R, Teixido-Tura G, Brion M, Forteza A, Rodriguez-Palomares J, Gutierrez L, et al. Relationship between fibrillin-1 genotype and severity of cardiovascular involvement in Marfan syndrome. Heart. 2017;103(22):1795–9. Epub 2017/05/05. doi: 10.1136/heartjnl-2016-310631. PubMed PMID: 28468757.

55. Odofin X, Houbby N, Hagana A, Nasser I, Ahmed A, Harky A. Thoracic aortic aneurysms in patients with heritable connective tissue disease. J Card Surg. 2021;36(3):1083–90. Epub 2021/01/22. doi: 10.1111/jocs.15340. PubMed PMID: 33476431.

56. Sermet-Gaudelus I, Boeck KD, Casimir GJ, Vermeulen F, Leal T, Mogenet A, et al. Ataluren (PTC124) induces cystic fibrosis transmembrane conductance regulator protein expression and activity in children with nonsense mutation cystic fibrosis. Am J Respir Crit Care Med. 2010;182(10):1262–72. Epub 2010/07/14. doi: 10.1164/rccm.201001-0137OC. PubMed PMID: 20622033.

57. Sermet-Gaudelus I, Renouil M, Fajac A, Bidou L, Parbaille B, Pierrot S, et al. In vitro prediction of stop-codon suppression by intravenous gentamicin in patients with cystic fibrosis: a pilot study. BMC Med. 2007;5:5. Epub 2007/03/31. doi: 10.1186/1741-7015-5-5. PubMed PMID: 17394637; PubMed Central PMCID: PMCPMC1852113.

58. Hamed SA. Drug evaluation: PTC-124--a potential treatment of cystic fibrosis and Duchenne muscular dystrophy. IDrugs. 2006;9(11):783–9. Epub 2006/11/11. PubMed PMID: 17096300.

59. Woodley DT, Cogan J, Hou Y, Lyu C, Marinkovich MP, Keene D, et al. Gentamicin induces functional type VII collagen in recessive dystrophic epidermolysis bullosa patients. J Clin Invest. 2017;127(8):3028–38. Epub 2017/07/12. doi: 10.1172/JCI92707. PubMed PMID: 28691931; PubMed Central PMCID: PMCPMC5531396.

60. Granata A, Serrano F, Bernard WG, McNamara M, Low L, Sastry P, et al. An iPSC-derived vascular model of Marfan syndrome identifies key mediators of smooth muscle cell death. Nat Genet. 2017;49(1):97–109. Epub 2016/11/29. doi: 10.1038/ng.3723. PubMed PMID: 27893734.

61. Davaapil H, Shetty DK, Sinha S. Aortic “Disease-in-a-Dish": Mechanistic Insights and Drug Development Using iPSC-Based Disease Modeling. Front Cell Dev Biol. 2020;8:550504. Epub 2020/11/17. doi: 10.3389/fcell.2020.550504. PubMed PMID: 33195187; PubMed Central PMCID: PMCPMC7655792.

62. Dosanjh A, Won CY. Amlexanox: A Novel Therapeutic for Atopic, Metabolic, and Inflammatory Disease. Yale J Biol Med. 2020;93(5):759–63. Epub 2021/01/01. PubMed PMID: 33380937; PubMed Central PMCID: PMCPMC7757066.

63. Hutchinson S, Furger A, Halliday D, Judge DP, Jefferson A, Dietz HC, et al. Allelic variation in normal human FBN1 expression in a family with Marfan syndrome: a potential modifier of phenotype? Hum Mol Genet. 2003;12(18):2269–76. Epub 2003/08/14. doi: 10.1093/hmg/ddg241. PubMed PMID: 12915484.

64. Aubart M, Gross MS, Hanna N, Zabot MT, Sznajder M, Detaint D, et al. The clinical presentation of Marfan syndrome is modulated by expression of wild-type FBN1 allele. Hum Mol Genet. 2015;24(10):2764–70. Epub 2015/02/06. doi: 10.1093/hmg/ddv037. PubMed PMID: 25652400.

65. Liao HK, Hatanaka F, Araoka T, Reddy P, Wu MZ, Sui Y, et al. In Vivo Target Gene Activation via CRISPR/Cas9-Mediated Trans-epigenetic Modulation. Cell. 2017;171(7):1495–507 e15. Epub 2017/12/12. doi: 10.1016/j.cell.2017.10.025. PubMed PMID: 29224783; PubMed Central PMCID: PMCPMC5732045.

66. Konermann S, Brigham MD, Trevino AE, Joung J, Abudayyeh OO, Barcena C, et al. Genome-scale transcriptional activation by an engineered CRISPR-Cas9 complex. Nature. 2015;517(7536):583–8. Epub 2014/12/11. doi: 10.1038/nature14136. PubMed PMID: 25494202; PubMed Central PMCID: PMCPMC4420636.

67. Sabatier L, Djokic J, Fagotto-Kaufmann C, Chen M, Annis DS, Mosher DF, et al. Complex contributions of fibronectin to initiation and maturation of microfibrils. Biochem J. 2013;456(2):283–95. Epub 2013/09/28. doi: 10.1042/BJ20130699. PubMed PMID: 24070235; PubMed Central PMCID: PMCPMC4550319.

68. Sheikh O, Yokota T. Developing DMD therapeutics: a review of the effectiveness of small molecules, stop-codon readthrough, dystrophin gene replacement, and exon-skipping therapies. Expert Opin Investig Drugs. 2021;30(2):167–76. Epub 2021/01/05. doi: 10.1080/13543784.2021.1868434. PubMed PMID: 33393390.

69. Finkel RS, Flanigan KM, Wong B, Bonnemann C, Sampson J, Sweeney HL, et al. Phase 2a study of ataluren-mediated dystrophin production in patients with nonsense mutation Duchenne muscular dystrophy. PloS one. 2013;8(12):e81302. Epub 2013/12/19. doi: 10.1371/journal.pone.0081302. PubMed PMID: 24349052; PubMed Central PMCID: PMCPMC3859499.

70. Nomakuchi TT, Rigo F, Aznarez I, Krainer AR. Antisense oligonucleotide-directed inhibition of nonsense-mediated mRNA decay. Nat Biotechnol. 2016;34(2):164–6. Epub 2015/12/15. doi: 10.1038/nbt.3427. PubMed PMID: 26655495; PubMed Central PMCID: PMCPMC4744113.

